# Monitoring single root hairs using a micro-chambered hydroponics system MiCHy reveals a new mode of action of the aminosteroid U73122

**DOI:** 10.1101/2023.07.20.549431

**Authors:** Hiromasa Shikata, Yoshikatsu Sato, Claus Schwechheimer

**Affiliations:** Division of Plant Environmental Responses, National Institute for Basic Biology, National Institutes of Natural Sciences, 38 Nishigonaka, Myodaiji, Okazaki, Japan; Course for Basic Biology, The Graduate University for Advanced Studies, SOKENDAI, Hayama, 240-0115, Japan; Chair of Plant Systems Biology, School of Life Sciences, Technical University of Munich, Emil-Ramann-Strasse 8, Freising, Germany; Institute of Transformative Bio-Molecules (WPI-ITbM), Nagoya University; Division of Biological Science, Graduate School of Science, Nagoya University

## Abstract

When using chemical probes and inhibitors for cell biological studies, the uniform and rapid access of the chemical to the sample is key for the accuracy of the analysis. Until now, however, this could only be accomplished by using expensive equipment and labor-intensive and often lengthy procedures. Here, we introduce MiCHy, a micro-chambered hydroponics system, as an accessible and versatile method for chemical treatments and demonstrate its use for imaging roots of *Arabidopsis thaliana*, the major model system for plant cell biological research. Using lipid biosensors in combination with established chemicals, such as the phosphoinositol 4-kinase (PI4K) inhibitor phenylarsine oxide and the phosphoinositide-specific phospholipase C (PI-PLC) inhibitor U73122, we demonstrate the suitability of MiCHy for studying the relationship between growth and lipid dynamics in root hairs. We further reveal novel effects of U73122 on the plasma membrane localization of phosphatidylinositol-4-phosphate 5-kinase 3 (PIP5K3), which is the first evidence demonstrating a role for U73122 besides its role as a PI-PLC inhibitor in plants.

## Introduction

Live-imaging of roots of the plant model organism *Arabidopsis thaliana* is a powerful tool to understand the cellular basis of cell growth, cell division, or responses to environmental stresses. Fluorescent proteins or dyes that visualize target biomolecules allow us to monitor their cellular distribution or cellular behavior. In standard imaging procedures, roots grown on solid or liquid media are directly transferred to imaging slides or soaked in the medium supplemented with chemicals like dyes or inhibitors before imaging by confocal microscopy and alike. Since roots sensitively respond to environmental changes in water availability, pH, nutrient concentration, osmolarity, and mechanical stimuli, such procedures can have unspecific effects that may be further enhanced during the time taken for sample preparation (Okada and Shimura, 1990; Takahashi, 1997; Verslues et al., 1998; Koyama et al., 2002; Potter et al., 2007). This asks for the development of fast and direct application and assay methods for chemical treatments of plant cells.

To minimize the undesired effects associated with mounting procedures, roots are often grown or acclimated on imaging slides or in cuvettes containing the growth media before imaging. Gösta Fåhraeus first described the methodology to stably monitor how root hairs respond to rhizobacteria in a micro-chamber, which was built on an imaging slide with a coverslip and contained a solid growth medium (Fåhraeus, 1957). The system, now known as the Fåhraeus slide, has subsequently been used and improved by many researchers to observe living root cells including root hairs (Supplemental Table S1). The use of liquid media in the Fåhraeus slide allows researchers to monitor root responses to treatments with chemicals such as inhibitors, dyes, or fixatives that can more readily reach the tissue and cells under investigation than when using solid media (Supplemental Table S1).

Microfluidics devices, where the medium can be changed during the experiment, represent another option to observe the living plant tissues and cells in various environmental conditions (reviewed by Yanagisawa et al., 2021). Microfluidics devices are made from transparent, gas-permeable silicone rubber, as a negative copy of the mold fabricated on a silicon wafer, and contain microchannels that are purpose-designed to grow roots to visualize root behavior. However, the fabrication of microfluidics devices and their use requires expensive equipment and perfusion systems, which limit their availability and access to a broad community.

Here, we describe a simple, cost-efficient system, micro-chambered hydroponics (MiCHy), for the stable monitoring of living roots and root hairs. The system is built from commercially available material and permits the study of chemical inhibitor effects on cells at a limited cost and without the necessity of a perfusion pump. We demonstrate the use of MiCHy for studies of the actions of dye staining and inhibitor treatments in intact roots and root hair cells. By imaging with MiCHy, we uncover that U73122, a well-known inhibitor of phosphoinositide-specific phospholipase C (PI-PLC), has another potential target in Arabidopsis root hairs.

## Results

### Design of MiCHy, a micro-chambered hydroponics system

Time lapse-imaging of roots and root hairs grown in solid media on imaging slides can minimize stress to plants associated with seedling transfer (and treatment), avoids root hair damage, and permits long-term imaging (Nakamura et al., 2018). However, when dyes or inhibitors are applied in such a system, treatment times and effects are dependent on their generally unknown diffusion rates in solid medium and possibly by their uneven penetration (Supplemental Movie S1).

To overcome these limitations, we developed an imaging system based on the Fåhraeus slides to visualize the living roots of *Arabidopsis thaliana* seedlings with simultaneous chemical treatments in a liquid growth medium (Fig. 1). In this device, an (approximately 2 cm × 2 cm) micro-chamber is built on a 2-well chambered coverglass. A thin space for root growth is sandwiched between the bottom coverglass and an upper hydrophilic plastic coverslip, and the interjacent growth space is made by small strips of a silicone rubber sheet on either side of the micro-chamber (Fig. 1A; Supplemental Fig. S1, A to G). In this device, *Arabidopsis thaliana* seeds can be positioned and germinated on the upper edge of the coverslip and grown in the vertically oriented micro-chamber, such that roots will grow in the space between the coverslip and the coverglass for microscopic observation (Fig. 1, A and B; Supplemental Fig. S1, H to N). The growth space is filled with a liquid growth medium and its thickness is adjustable by adjusting the thickness of the silicone rubber strips (0.3 to 1 mm) (Fig. 1B; Supplemental Fig. S1J).

**Figure 1.**
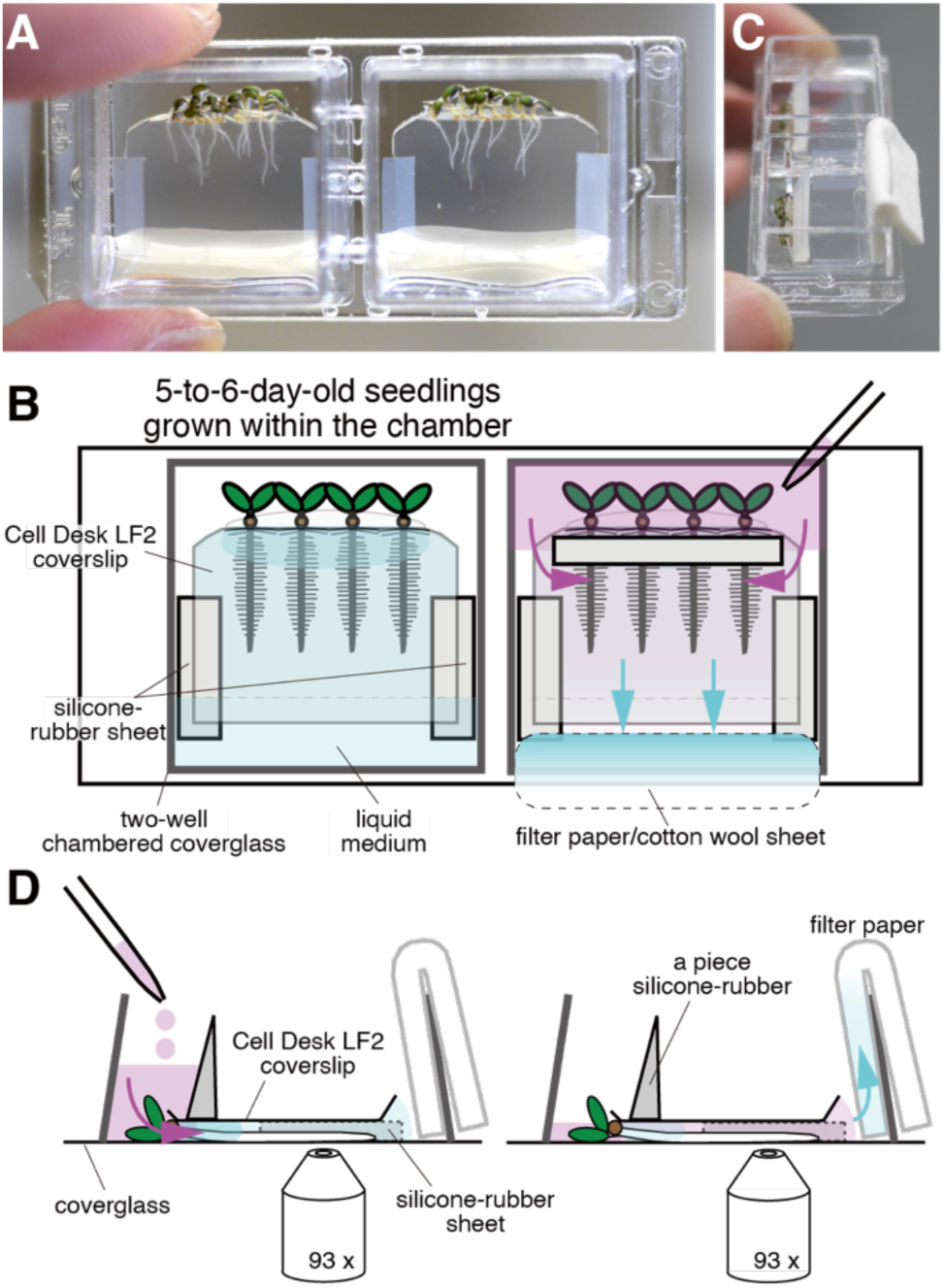
Design of the micro-chambered hydroponics system MiCHy. **A)** A representative image of 5-day-old seedlings grown in the micro-chamber. Roots are grown in a thin space between a bottom coverglass and an upper Cell Desk LF2 coverslip, filled with the liquid medium. **B to D)** Schematic illustrations on the structure of the micro-chamber and procedure for medium exchange. Arrows indicate the flow of the growth media when it is loaded on the upper part of the micro-chamber with a pipet. A sheet of filter paper or cotton wool is fitted in the lower gap between the coverslip and plastic chamber to efficiently exchange medium as shown in **C**) to **D**). The micro-chamber is mounted on the inverted microscope and image acquisition and medium exchange can be performed coincidentally.

When establishing this system, we initially used glass coverslips for the upper layer as used in the Fåhraeues slides (Fig. 1) (Fåhraeues, 1957). When using glass coverslips, we found, however, that roots grew much shorter and had fewer root hairs when growing in the center of the chamber than when grown at the sides of the chamber (Supplemental Fig. S2, A to C). Since silicone has a high gas permeability, we assumed that low oxygen availability in the center of the micro-chamber may affect root and root hair growth. When we chose a hydrophilic, biocompatible plastic coverslip Cell Desk LF2 with expecting improved gas permeability in the micro-chamber, we indeed obtained uniform root and root hair growth in the entire micro-chamber (Supplemental Fig. S1, A, D, and E). We obtained similar results when combining a gas-permeable plastic bottom dish (e.g. Ibidi μ-slide 2 well-chambered coverslip) with a glass coverslip as the upper layer.

To measure oxygen diffusion within the glass coverslip- or Cell Desk LF2-layered micro-chambers, we performed colorimetric oxygen detection assays (Barrett-Lennard and Dracup, 1988). Based on the rapid coloration of the assay, we confirmed high oxygen diffusion from the rubber sheet to the medium as well as from the surrounding air, which was detectable in 4 mins (Supplemental Fig. S2F, Movie S2). In contrast, coloration inside the Cell Desk LF2-layered chamber was comparatively delayed (80 mins), but even more so when using the glass coverslip (180 mins). Oxygen diffusion thus correlated with the root growth differences observed in the two types of micro-chambers (Supplemental Fig. S2, A to E). We decided to use the Cell Desk LF2 as the upper coverslip for our further experiments.

The growth medium in the device is exchangeable by manual loading with a pipet, even during imaging on inverted microscopes. For smooth loading and exchange of the medium, we stuck a small silicone-rubber piece on the upper part of the plastic coverslip and fitted a sheet of filter paper or cotton wool on the lower gap between the coverslip and plastic chamber (Fig. 1, B and C). The medium can be supplemented with chemicals of choice such as inhibitors or dyes and applied to the upper part of the device with a pipet (Fig. 1, B and C). The medium in the chamber can be readily replaced with new medium and buffer exchanges for washing-out treatments are carried out in the same manner (Supplemental Movies S3 and S4). We have named this <u>mi</u>cro-<u>c</u>hambered <u>hy</u>droponics system MiCHy.

### Staining living roots with MiCHy

Dye staining of living roots provides insights into how biomolecules or organelles behave in cells. Arabidopsis roots are generally stained after transferring seedlings grown on a solid medium with forceps into a liquid medium, supplemented with the respective dye. During this procedure, fragile tissues like root hairs are easily damaged by mechanical or osmotic stress. Other issues associated with this comparatively lengthy procedure are the diffusion and translocation of dyes into tissues and cells after prolonged incubation or their secondary effects on cellular processes.

The dye FM4-64X is rapidly taken up by the plasma membrane within a few minutes and translocated into the inner membrane by endocytosis (Ovečka et al., 2005). Conventional mounting procedures, which take comparatively long, are too slow to allow imaging of this fast process. We used 2 µM FM4-64X in MiCHy and successfully visualized the plasma membranes of root hairs and epidermal cells already one min after the application of FM4-64X (Supplemental Movie S5). The analysis revealed a rapid decrease of the plasma membrane signal, which reduced faster in root hairs than in epidermal cells, which may reflect a difference in the endocytosis rates between the two cell types.

Next, we stained roots with the cell wall pectin stain propidium iodide (PI) (Rounds et al., 2011). With MiCHy, the cell wall was visualized in one min after 10 µM PI application and its signal intensity increased during the observation (Supplemental Movie S6). We also applied the sterol stain Filipin III (10 µg/ml), which is known to stain the apical plasma membrane and globular particles in the cytosol of the growing root hair tip, and to inhibit root hair growth or to stain the entire plasma membrane in non-growing root hairs (Ovečka et al., 2010). We found that the apical plasma membrane of growing root hairs was stained with Filipin III, which is accompanied by cell growth arrest, 30 sec after application, and that the fluorescent signal gradually spread throughout the plasma membrane and internalized into the cytosol (Supplemental Movie S7). We also detected enlarging Filipin III-stained particles, which emerged on the plasma membrane at the root hair tip approximately 2 min after the application, suggesting that these particles represent the aggregates of endocytic vesicles that include Filipin-sterol complexes (Supplemental Movie S7), as described previously (Ovečka et al., 2010). These results indicate that MiCHy is useful for the dye staining of living cells, including staining with diffusive and toxic dyes.

### Brefeldin A treatment of living roots

To evaluate the use of MiCHy for treatments with inhibitory chemicals, we grew Arabidopsis seedlings overexpressing YFP-fused D6PK in MiCHy (Zourelidou et al., 2009). YFP-D6PK is a polarly localized protein kinase at the basal plasma membrane of root cells and its intracellular trafficking is sensitive to Brefeldin A (BFA), an inhibitor of the ARF-GEF GNOM (Barbosa et al., 2014). We conducted time-lapse imaging of YFP-D6PK before and after treatment with 50 µM BFA. YFP-D6PK was observed at the basal plasma membrane before the treatment, visibly internalized to the cytosol already 3 min after the treatment, and almost fully internalized after 10 min (Supplemental Movie S8). Subsequent BFA wash-out treatments with growth medium recovered the localization at the basal plasma membrane already detectably after 3 min and almost fully after 10 min (Supplemental Movie S8). This behavior, consistent with our previous work (Barbosa et al., 2014), proved the usefulness of this system for chemical treatments of roots and dynamics analyses of cell biological processes.

### Phospholipid dynamics in root hairs treated with phenylarsine oxide (PAO) and U73122

We next analyzed the dynamics of the biosensors for phosphatidylinositol 4-phosphate [PtdIns(4)P] and PtdIns(4,5)P_2_ in treatments of their biosynthesis or metabolic inhibitors in growing root hairs (Fig. 2A). Phenylarsine oxide (PAO) inhibits PI4 kinase, which produces PtdIns(4)P from PtdIns, and the aminosteroid U73122 inhibits PI-PLC, which converts PtdIns(4,5)P_2_ into diacylglycerol (DAG) and Ins(1,4,5)P_3_ (IP_3_) (Bleasdale et al., 1989; Balla et al., 2002).

**Figure 2.**
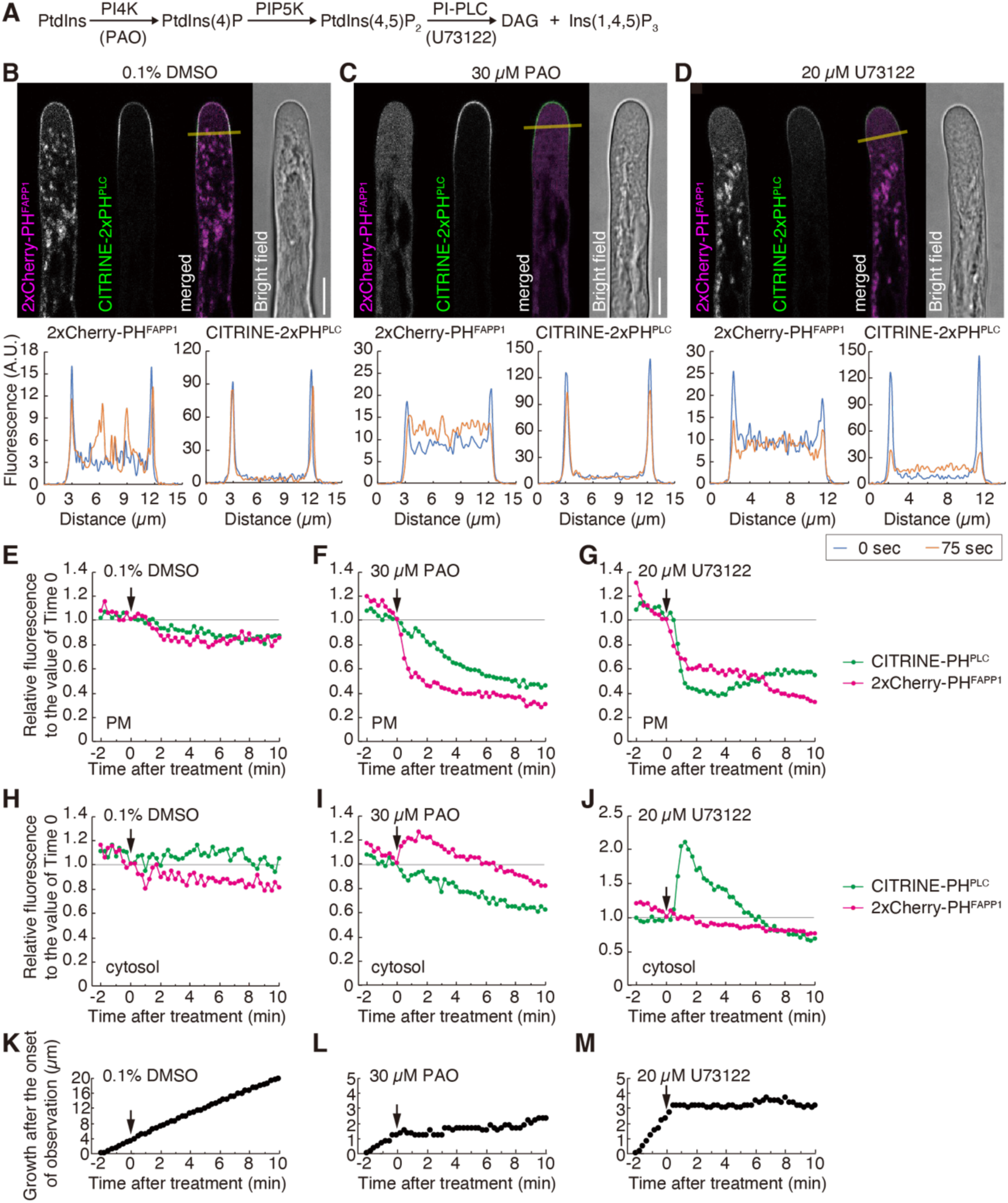
Live-imaging of the PtdIns(4)P and PtdIns(4,5)P_2_ biosensors in MiCHy. **A)** Schematic of the pathway of biosynthesis and metabolism for PtdIns(4)P and PtdIns(4,5)P_2_. Chemicals used in this study, which are known to inhibit biosynthetic steps (arrows), are shown in parentheses placed below the arrows. Root hairs expressing PtdIns(4)P biosensor 2xCherry-PH^FAPP1^ and PtdIns(4,5)P_2_ biosensor CITRINE-PH^PLC^ were treated with 0.1% DMSO as control (**B, E, H, K**), 30 µM PAO (**C, F, I, L**), or 20 µM U73122 (**D, G, J, M**). Representative images of a root hair 75 sec after treatment with each drug was shown in **B**) to **D**). The bottom graphs present the distribution of fluorescence signals on the line of interest (shown as yellow lines) in the images at 0 and 75 sec after the treatment. White bars represent 10 µm. Changes in fluorescence intensities on the plasma membrane (PM) or in the cytosol at the root hair tips were quantified as described in Supplemental Fig. S4, and the relative values to that of Time 0 are plotted in **E**) to **G**) or **H**) to **J**), respectively. In **K**) to **M**), the graphs show root hair elongation after the onset of observation. Arrows indicate the time when the chemicals were applied. The experiments were repeated at least three times and the representative results are shown.

The PtdIns(4)P biosensor 2xCherry-PH^FAPP1^ is distributed on the plasma membrane predominantly at the tips of root hairs and in endomembranes like the Golgi apparatus, while the PtdIns(4,5)P_2_ biosensor CITRINE-2xPH^PLC^ is only observed on the apical plasma membrane (Fig. 2, B, E, and H; Supplemental Movie S9) (Van Leeuwen et al., 2007; Thole et al., 2008). We generated Arabidopsis plants expressing the PtdIns(4)P and PtdIns(4,5)P_2_ biosensors and treated the roots with PAO or U73122. 30 µM PAO treatment inhibited, within ∼1 min of treatment, root hair elongation accompanied by an arrest of cytoplasmic streaming, which correlated with complete depletion of the PtdIns(4)P biosensor from both the plasma membrane and endomembranes, at the expense of an increase in cytosolic signal (Fig. 2, C, F, I, and L; Supplemental Movies S10). This result is consistent with the fact that PI4 kinases are localized to the trans-Golgi network and the plasma membrane (Kang et al., 2011; Noack et al., 2022).

Root hair growth was arrested immediately after treatment with 5 and 20 µM U73122, detectable as early as ∼1 min after the treatment. In turn, cytoplasmic streaming was gradually reduced at both concentrations, which distinguished U73122 from PAO treatments (Fig. 2, D, G, J, and M; Supplemental Movies S11 and S12). Unexpectedly, U73122 treatment led to a detectable internalization of the PtdIns(4,5)P_2_ biosensor CITRINE-2xPH^PLC^ in one min after the treatment, but a little of the biosensor remained detectable at the plasma membrane throughout the observation (∼10 min). 2xCherry-PH^FAPP1^, the biosensor for PtdIns(4,5)P_2_-precursor PtdIns(4)P, was also visibly decreased on the plasma membrane after U73122 treatment, however, without a detectable internalization, and the reduction of CITRINE-2xPH^PLC^ on the plasma membrane was faster than that of 2xCherry-PH^FAPP1^ (Fig. 2, G and J). These suggest that the decreased 2xCherry-PH^FAPP1^ signal may be a secondary effect of the disturbed PtdIns(4,5)P_2_ level or reflect technical issues such as photo-bleaching, which could be enhanced by a decreased turnover of the protein as a consequence of arrested cytoplasmic streaming (Fig. 2).

To test whether the U73122-induced reduction of the PtdIns(4,5)P_2_ biosensor on the plasma membrane resulted from PI-PLC inhibition, we applied the PI-PLC inhibitor neomycin, which binds negatively charged phospholipids, including PtdIns(4,5)P_2_, and inhibits their biological activity (Zhao et al., 2004). We observed arrested root hair growth and transient reduction of the PtdIns(4,5)P_2_ biosensor on the plasma membrane in neomycin treatment (Supplemental Fig. S3, Movie S13). However, the internalization was mild and transient when compared to U73122 treatment, and the signal at the plasma membrane remained unevenly but strongly on the plasma membrane until an arrest of cytoplasmic streaming (Supplemental Fig. S3, Movies S10 to S13). These results suggest that U73122 and neomycin have distinct effects on PtdIns(4,5)P_2_ turnover at the concentrations used in our study.

### The turnover of diacylglycerol during U73122 treatment

To rationalize the above-described U73122-induced depletion of PtdIns(4,5)P_2_, we considered two possible scenarios. First, U73122 could activate, rather than inhibit, PI-PLC in root hairs, as it had previously been reported for U73122 in human cells (Klein et al., 2011). Second, U73122 could affect the biosynthesis of PtdIns(4,5)P_2_. To get further insight into the above-described U73122-induced depletion of PtdIns(4,5)P_2_ on the plasma membrane, we analyzed the accumulation of DAG, the product of PtdIns(4,5)P_2_ hydrolysis by PI-PLC, after U73122 treatment. To this end, we generated Arabidopsis plants expressing the DAG biosensor 2xC1a^PKCγ^ fused to YFP (YFP-2xC1a^PKCγ^) (Oancea et al., 1998), under the control of the root hair-specific *EXP7* promoter (Cho and Cosgrove, 2002). We confirmed the previously described plasma membrane localization of YFP-2xC1a^PKCγ^ in growing root hairs (Fig. 3) (Vermeer et al., 2017). We further found that YFP-2xC1a^PKCγ^ was internalized from the plasma membrane immediately after U73122 treatment, in the same manner as the PtdIns(4,5)P_2_ biosensor (Fig. 3; Supplemental Movie S14), suggesting that PI-PLC was not activated after U73122 treatment. At the same time, neomycin treatment did not affect YFP-2xC1a^PKCγ^ accumulation on the plasma membrane at root hair tips (Fig. 3). This suggests that DAG levels are relatively stable on the plasma membrane even when the production of DAG by PI-PLC is inhibited. Thus, we assumed that the U73122-induced reduction of DAG on the plasma membrane could reflect the depletion of its precursor PtdIns(4,5)P_2_ on the plasma membrane.

**Figure 3.**
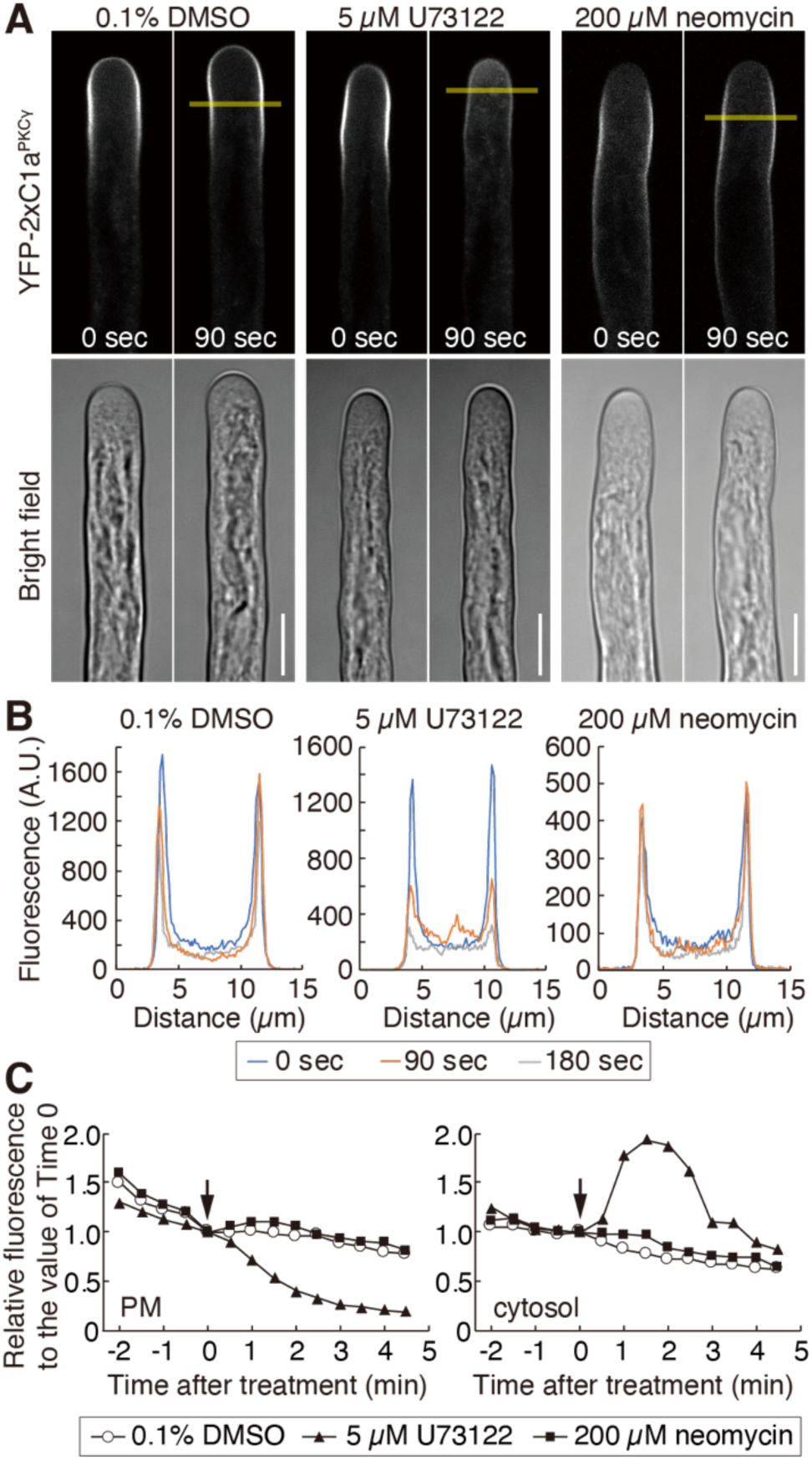
Live-imaging of the DAG biosensor responding to inhibitors in MiCHy. Root hairs expressing the DAG biosensor YFP-2xC1a^PKCγ^ were treated with 5 µM U73122, 200 µM neomycin, or 0.1% DMSO as control. **A)** Representative images of a root hair at 0 and 90 sec after treatment of each chemical are shown. White bars represent 10 µm. **B)** Graphs representing the distribution of fluorescence signals on the line of interest (LOI) in the images at 0, 90, and 180 sec after the treatment. LOIs are shown as yellow lines in **A**. **C)** Changes in fluorescence intensities on the plasma membrane (PM) or in the cytosol at the root hair tips were quantified as described in Supplemental Fig. S4, and the relative values to that of Time 0 are plotted. Arrows indicate the time when the chemicals were applied. The experiments were repeated at least three times and the representative results are shown.

### Localization of PIP5K3, a biosynthesis enzyme for PtdIns(4,5)P_2_, during U73122 treatment

To test whether U73122 affected the biosynthesis of PtdIns(4,5)P_2_, we analyzed the effect of U73122 on the subcellular localization of YFP-fused PtdIns-4-phosphate 5-kinase 3 (PIP5K3-YFP), which is a major enzyme to convert PtdIns(4)P to PtdIns(4,5)P_2_ in root hairs. PIP5K3-YFP was localized to the apical plasma membrane of growing root hairs as reported previously (Fig. 4, A and B; Supplemental Movie S15) (Kusano et al., 2008; Stenzel et al., 2008). We found, however, that U73122 treatment immediately diminished the localization of PIP5K3-YFP on the plasma membrane, which was similar to the behavior of the PtdIns(4,5)P_2_ biosensor (Figs. 2D and 4; Supplemental Movie S15). By contrast, neomycin treatment affected no change in the localization pattern of PIP5K3-YFP (Fig. 4, A and B). Those findings indicate that the U73122-induced depletion of PtdIns(4,5)P_2_ may result from reduced PIP5K3 levels at the apical plasma membrane (Fig. 4D).

**Figure 4.**
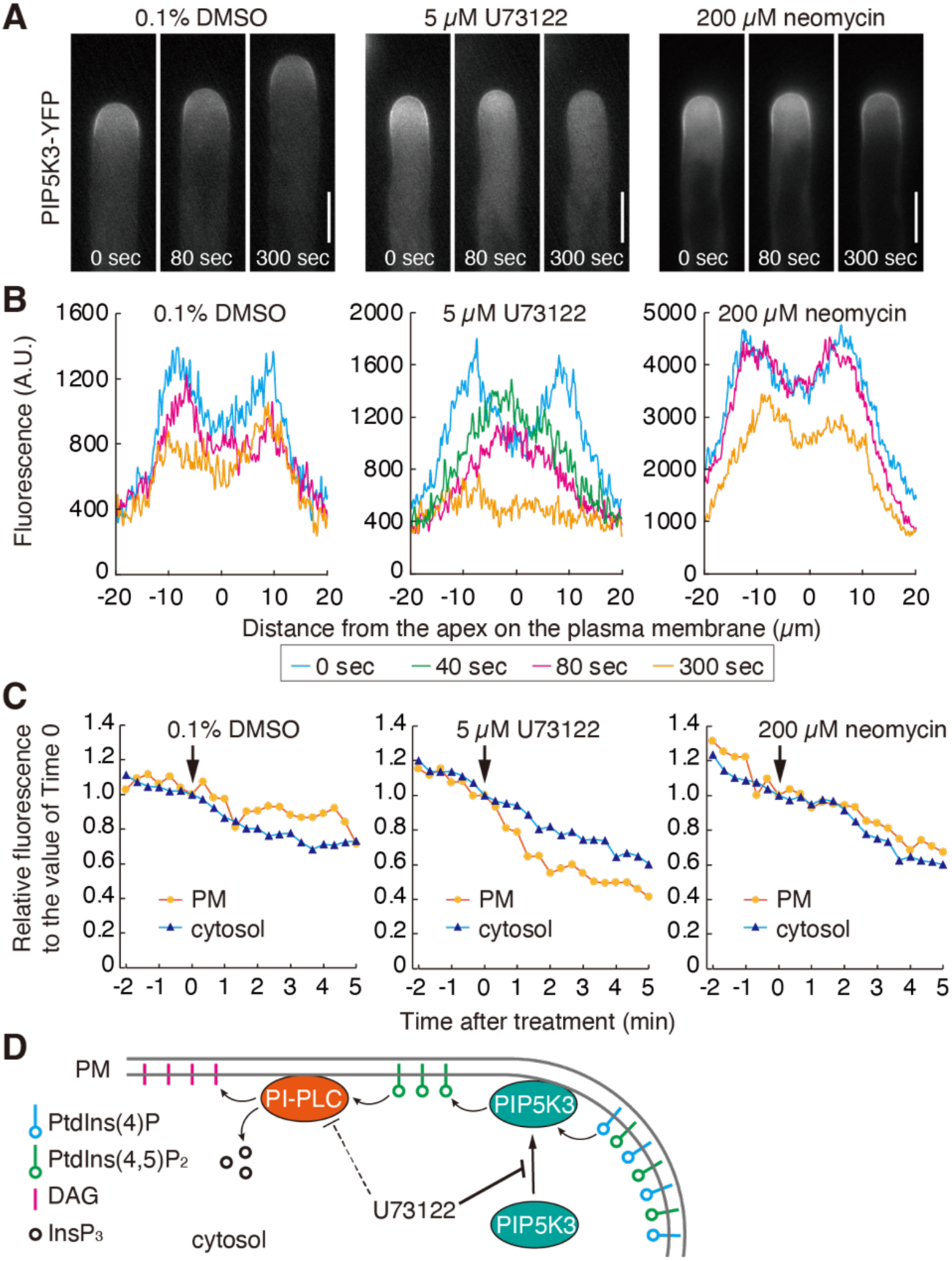
Live-imaging of PIP5K3 responding to inhibitors in MiCHy. Root hairs expressing PIP5K3-YFP were treated with 5 µM U73122, 200 µM neomycin, or 0.1% DMSO as control. **A)** Representative images of root hairs at 0, 80, or 300 sec after treatment of each chemical. White bars represent 10 µm. **B)** Graphs representing the distribution of fluorescence signals on the plasma membrane at the root hair tips at 0, 40, 80, and 300 sec after the treatment in the images shown in **A**). The point 0 in the x-axis indicates the position of a root hair apex as described in Supplemental Fig. S4. **C)** Changes in fluorescence intensities on the plasma membrane (PM) or in the cytosol at the root hair tips were quantified, and the relative values to that of Time 0 are plotted. Arrows indicate the time when the chemicals were applied. The experiments were repeated at least three times and the representative results are shown. **D)** Schematic illustration of the modes of action of U73122 in the growing root hair tip. U73122 may inhibit PIP5K3 translocation from the cytosol to the plasma membrane, thereby leading to the reduction of PtdIns(4,5)P_2_ and the resulting reduction of DAG on the plasma membrane of root hair tips.

## Discussion

Fåhraeus slides and their derivative micro-chambers are useful tools to visualize living roots and root hairs. However, as our study demonstrated, limited oxygen availability in these chambers negatively affects root growth, and we introduce MiCHy as a related system with improved oxygen permeability (Supplemental Fig. S2). Indeed, previous work showed that low oxygen conditions reduce root growth and root hair number (Loehwing, 1934; Durell, 1941). Ketelaar et al. (2004) later reported an improved method, in which roots were grown on the solid medium on an imaging slide and were covered with a gas-permeable film to observe living root hairs. While this method enables us to treat intact roots with chemicals, the chemical diffusion is comparatively slow (> 10 min) and reaches the samples in the solid medium with an unpredictable delay (Ketelaar, 2014), as also demonstrated by us here using a similar system (Supplemental Movie S1). We introduce the use of oxygen-permeable materials, namely silicone rubber sheets and a Cell Desk LF2 coverslip (or related), which allows us to monitor intact and healthy roots and root hairs before and after chemical treatments with a high temporal resolution for reliable and reproducible imaging.

Using MiCHy, we demonstrate that treatments with the well-known PI-PLC inhibitor U73122, decrease the level of the PLC-substrate PtdIns(4,5)P_2_ on the plasma membrane in growing root hairs. This observation is opposite to the expected effect that U73122 inhibits PI-PLC activity, thereby increasing the amount of PtdIns(4,5)P_2_ on the plasma membrane (Staxén et al., 1998; Van Leeuwen et al., 2007). In mammals, U73122 has been claimed not to be specific to PI-PLC, and the situation in plants is predicted to be the same as in mammals, though no evidence has been reported so far (Munnik 2014). Hence, we provide, as far as we are aware, the first support for this type of effect of U73122 in plants. Since it had previously been reported that recombinant PI-PLC from *Commelina communis* is not inhibited by 5 µM of U73122 (Staxén et al., 1998), the concentration used in our study, we suggest that the effects of U73122 observed by us may be independent of PI-PLC and represent a new mode of action in plants. Along these lines, we would also like to note that the effects of U73122 could be observed within minutes whereas previous studies had used hourlong incubations (Figs. 2 to 4) (Staxén et al., 1998; Van Leeuwen et al., 2007). Thus, we cannot rule out the possibility that U73122 inhibits PI-PLC after prolonged treatments in *Arabidopsis thaliana*.

The U73122-induced reduction of PIP5K3 levels on the plasma membrane could account for the depletion of PtdIns(4,5)P_2_. In contrast, we found, however, that U73122 treatment also reduced PtdIns(4)P levels (Fig. 2). We consider it unlikely that the depletion of PtdIns(4,5)P_2_ on the plasma membrane directly resulted from the reduction of PtdIns(4)P level because the extent of PtdIns(4,5)P_2_ reduction was comparatively mild in the PAO treatment, which completely diminished the PtdIns(4)P level on the plasma membrane and endomembrane (Fig. 2). Noack et al. (2022) speculated that a PI4Kα1 complex, which produces PtdIns(4)P on the plasma membrane, could be recruited to the plasma membrane domain in which anionic membrane lipids, such as PtdIns(4,5)P_2_, are enriched. Given that hypothesis, reduced levels of PtdIns(4)P after U73122 treatment could be explained as a consequence of the depletion of PtdIns(4,5)P_2_ level on the apical plasma membrane.

After PAO treatment, the levels of PtdIns(4,5)P_2_ and DAG were relatively stable on the plasma membrane regardless of the decreased supply of their precursor PtdIns(4)P (Figs. 2 and 3), implying that PAO also might affect the metabolism of PtdIns(4,5)P_2_ and DAG. As PI-PLC requires Ca^2+^ for its activation (Drøbak, 1992), we cannot dismiss the possibility that PAO could decrease cytosolic Ca^2+^ concentration in root hairs, thereby reducing the activity of PI-PLC in the cells. Further experiments on Ca^2+^ imaging will be helpful to explain this mode of PAO action.

U73122 affects the plasma membrane localization of PIP5K3, which belongs to the type-A subgroup of PIP5Ks containing MORN motifs that could serve to associate the proteins to the plasma membrane (Fig. 4) (Mueller-Roeber and Pical, 2002). How PIP5K3 is localized to the plasma membrane remains unknown but it has been reported that the plasma membrane-associated proteins BREVIS RADIX (BRX) and PROTEIN KINASE ASSOCIATED WITH BRX (PAX) recruit the PIP5K3-related and partially redundant PIP5K1/2 to the plasma membrane in the protophloem (Marhava et al., 2020; Watari et al., 2022). A related mechanism may thus exist for PIP5K3 recruitment. During root hair initiation, several proteins, including PIP5K3, are accumulated at a sterol-enriched domain (Stanislas et al., 2015). Hence, membrane lipids or plasma membrane-associated proteins, which could be required for the PIP5K3 localization, are potential targets for U73122. Further studies will need to seek the actual targets of U73122 in root hairs.

## Conclusion

This study introduced the micro-chambered hydroponics system MiCHy as an accessible tool to visualize roots with simultaneous chemical treatments and characterized the mode of action of chemicals such as well-known probes and inhibitors with a high temporal resolution. The use of this system can minimize the mechanical or osmotic stresses to plants, which could arise in standard imaging procedures, and allows us to monitor intact, identical cell(s) or tissue(s) before and after chemical treatments. The advantage of this system is its simple structure that can be assembled from commercially available and affordable materials without the costly need for a perfusion pump. MiCHy will give a broad plant research community the opportunity to expand their research for cell biological analyses in roots by using this cost-efficient method.

## Materials and Methods

### Plant materials

All experiments performed in this study used wild-type *Arabidopsis thaliana* (L.) Heynh Columbia-0 (Col-0) or transgenic lines expressing the reporters P5R (2xCherry-1xPH^FAPP1^) for the detection of PtdIns(4)P), P24Y (CITRINE-2xPH^PLC^) for the detection of PtdIns(4,5)P_2_) (Simon et al., 2014), *35Sp::YFP-D6PK* (Zourelidou et al., 2009), or *PIP5K3p::PIP5K3-YFP* (line #4) (Kusano et al., 2004). The line expressing both, 2x Cherry-1xPH^FAPP1^ and CITRINE-2xPH^PLC^ was obtained by crossing.

### Plasmid construction

For the cloning of *EXP7p::YFP-2xC1a^PKCγ^*, the *YFP* coding sequence was amplified by overlap extension PCR (Supplemental Table S2) to mutate an internal *Aau*I site, and the product was digested with *EcoR*I and *Aau*I. That DNA fragment, as well as an *Aau*I/*Not*I-digested fragment including the *CaMV 35S* terminator (35St) obtained from the pExtag vector (Zourelidou et al., 2009), were inserted into the vector pGreenII0179 and the resulting vectors were designated pGreen0179_YFP-(*Aau*I)::35St. For expression in root hairs, a 725-bp *EXPANSIN7A* promoter fragment (*EXP7*p), amplified by PCR from *Arabidopsis thaliana* genomic DNA, was inserted into the *Kpn*I and *Xho*I sites of pGreen0179_YFP-(*Aau*I)::35St. Two tandem copies of the cysteine-rich 1a (C1a) domain (amino acids 26-86) from rat PKC gamma, codon-optimized for *Arabidopsis thaliana* and fused with a GGGSGGG linker, were chemically synthesized (Thermo Fisher Scientific, MA, USA) (Supplemental Table S2). The DNA fragment was introduced into pGreen0179_YFP-(*Aau*I)::35St with *EXP7a* promoter by using a NEBuilder HiFi DNA Assembly kit (NEB, MA, USA), and the resulting construct was transformed into Arabidopsis Col-0 by floral dip (Clough and Bent, 1998).

### Preparation of micro-chambers

The micro-chamber was constructed on the 2-well chambered coverglass (Nunc^®^ Lab-Tek^TM^; Thermo Fisher Scientific) (Supplemental Fig. S1). Other commercially available chambered coverglasses from Iwaki (Shizuoka, Japan) and Eppendorf (Hamburg, Germany) are applicable as well. Two slips of the silicone rubber sheet (4 mm × 10 mm, 0.3 - 1.0 mm thick; AS ONE, Osaka, Japan) were positioned on the surface of a bottom glass in one well (Supplemental Fig. S1F) and a glass coverslip or Cell Desk LF2 coverslip (Sumitomo Bakelite, Tokyo, Japan), cut with a surgical knife and folded (Supplemental Fig. S1, A to D) was stuck on top of the silicone rubber slips to generate a gas permeable growth chamber (Supplemental Fig. S1G). For our studies, we routinely used 0.3-mm- or 0.5-mm-thick silicone rubber sheets for imaging of main roots or root hairs, respectively. When using the 0.3-mm-thick sheet, Arabidopsis seed could be directly positioned with the help of a pipet in the gap between the coverslip and the bottom glass. When silicone rubber sheets with 0.5-mm-thickness or thicker were used, a thin layer of melted 0.8% GTG-Agarose (Lonza, Basel, Switzerland) was poured into the upper space between the bottom glass and the coverslip to allow positioning of the seeds (Supplemental Fig. S1H).

To grow the seedlings in the liquid medium, we applied 500 µL of ½ MS medium (2.21 g/l Murashige & Skoog basal salts including Gamborg B5 vitamins [Duchefa Biochemie, Haarlem, The Netherlands]), 0.5 g/l 2-N-morpholino ethane-sulfonic acid [MES], pH 5.8) supplemented with 2% sucrose on the bottom part of the micro-chamber with a pipet (Supplemental Fig. S1J). The chamber was covered with its lid and fixed with surgical tape (3M, Maplewood, MN, USA), and kept vertically at an ∼70° angle in a transparent humid container under continuous white light (∼80 µmol/m^2^/s) for 5 - 6 days at 22°C before further use (Supplemental Fig. S1, K to N).

### Fluorescent dye staining and chemical treatments

One day before the onset of treatments, the growth medium in the micro-chamber was replaced with fresh ½ MS medium to reduce stress during subsequent chemical treatments. For observation, the extra liquid medium was removed from the micro-chamber and then a silicone-rubber piece and a sheet of filter paper or cotton wool were placed on the coverslip or the bottom of the chamber, respectively. The micro-chamber was set on the inverted microscope, as specified below, and roots that came out of the agar bed, if used, were observed, since diffusion of chemicals into agar is slower than into liquid. Before image acquisition, 150 µL of the liquid media was applied at the top of the chamber with a pipet, and root hairs that remained immobile during the liquid application were chosen for imaging. After the onset of time-lapse observations, 300 µL of liquid media supplemented with inhibitors or fluorescent dyes were applied with a pipet to the top of the chamber. For wash-out treatments, 500 µL of the growth medium was applied from the top of the chamber twice. The following inhibitors were dissolved in DMSO and used at final concentrations of 30 µM PAO (Sigma Aldrich, Taufkirchen, Germany), 5 or 20 µM U73122 (Sigma Aldrich), and 10 µg/mL Filipin III (Santa Cruz Biotechnology, Dallas, TX, USA). 0.1% DMSO was used as a solvent control. Neomycin (Wako Chemicals, Tokyo, Japan), PI (propidium iodide; Sigma Aldrich), and FM4-64X (Invitrogen, Carlsbad, CA, USA) were dissolved in water and used at 200 µM, 10 µM, and 2 µM final concentration, respectively.

### Image Acquisition

Root hairs expressing 2xCherry-PH^FAPP1^ and CITRINE-2xPH^PLC^, as well as roots expressing YFP-D6PK, or PI- or FM4-64-stained roots were observed with a Leica SP8 FALCON inverted confocal microscope with HyD detectors (Leica Microsystems, Jena, Germany). A glycerol-immersion 93x objective lens with motorized correction collar (HC PL APO 93x/1.30 GLYC motC STED W; Leica Microsystems), or a dry 10x or 20x objective lens (HC PL APO CS2 10x/0.40 DRY or 20x/0.75 DRY, respectively; Leica Microsystems) were used for imaging of root hairs or main roots, respectively. CITRINE and YFP were excited with a 514-nm laser and the fluorescent wavelength range from 525 to 570 nm was detected. FM4-64X or PI were excited with 514 or 488 nm laser, respectively, and the fluorescent wavelength ranges from 650 to 750 nm or 600 to 660 nm were detected, respectively.

Root hairs expressing CITRINE-2xPH^PLC^ or YFP-2xC1a^PKCγ^ were observed with an FV1000 inverted confocal microscope (Olympus, Tokyo, Japan) with a water-immersion 60x objective (UplanApo 60x/IR, NA1.2; Olympus). CITRINE and YFP were excited with a 514 nm laser and the fluorescent wavelength range from 530 to 630 nm was detected.

Filipin III-stained root hairs and root hairs expressing PIP5K-YFP were observed with an inverted epifluorescence microscope (Eclipse Ti2-E; Nikon, Tokyo, Japan) equipped with a CMOS camera (ORCA-Fusion BT; Hamamatsu Photonics, Hamamatsu, Japan) and a silicone-immersion 100x objective lens (SR HP Plan Apo Lambda S 100XC Sil, NA, 1.35; Nikon). Fluorescence from YFP or Filipin III was detected with the filter sets YFP-2427B-NTE (Sermrock, Rochester, USA) or U-MWU (Olympus), respectively. The excitation light for YFP was passed through the long-pass filter FF01-496/LP-25 (Sermrock) to cut off wavelengths below 501 nm to minimize the autofluorescence of root hairs.

### Image quantification

To quantify the dynamics of the biosensors and PIP5K3-YFP during treatments, we measured fluorescence intensities on the plasma membrane around the apex of root hairs and in the cytosol of root hair tips. The apex of a root hair was determined as an intersection between the plasma membrane and an extension of a skeletonized line produced from the binary image, which was processed from a fluorescence image with the ImageJ software (https://imagej.nih.gov/ij/) (Supplemental Fig. S4, A to F). 0.6-µm- or 0.5-µm-thick best-focused slices were selected manually from Z-stack images with 0.3 or 0.5 µm intervals, respectively, and were converted into a z-axis sum projection image. A line of interest (LOI) was drawn on the plasma membrane around the apex and the intensity profile in the LOI was obtained (Supplemental Fig. S4, G and H). The intensities in a range from -20 µm to +20 µm were summarized as the intensities on the plasma membrane, except for PIP5K3-YFP. For PIP5K3-YFP, intensities from -15 to +15 µm were summed up. All image processing and measurements were performed using the ImageJ software. For quantification of the images acquired with the epifluorescence microscope Nikon Ti-2E, background signals were subtracted by the “subtraction background” algorithm of the ImageJ software.

### Detection of oxygen diffusion

Relative rates of oxygen diffusion into the micro-chambered hydroponics were visualized by the colorimetric test described by Barrett-Lennard and Dracup (1988). Sets of slips of a 0.5 mm-thick silicone rubber sheet and a glass coverslip (18 mm ×18 mm) or a shaped Cell Desk LF2 coverslip were stuck onto a well-bottom of the two-well chambered coverglass. Degassed water containing 5 mM Na_2_S_2_O_4_ (Wako Chemicals), 0.1 mM methylene blue (Wako Chemicals), and 0.8% (w/v) GTG-Agarose was injected into the space between the well-bottom and the coverslips, and image acquisition was started after the onset of injection and take images in a 4-min interval. Atmospheric oxygen diffused into the reaction colors it deep blue.

## Acknowledgments

We are grateful to Drs. Markus Grebe (University of Potsdam, Germany), Moritaka Nakamura (formerly at the Universities of Potsdam, Nagoya, Japan, and National Institute for Basic Biology (NIBB), Japan), and Siori Nagahara (Nagoya University, Japan) for sharing their plant imaging techniques, and Dr. Kagayaki Kato (NIBB) for the helpful advice on image quantification. *PIP5K3p::PIP5K3-YFP* seeds were a kind gift from Dr. Takashi Aoyama (Kyoto University, Japan). Confocal microscopy imaging was performed at the ITbM Live Imaging Center, Nagoya University, and the Division of Plant Environmental Responses, NIBB. The support of plant cultivation rooms was provided by the Model Plant Research Facility of NIBB. We thank Mayako Hamada and Yoriko Soma (NIBB) for technical assistance and the Arabidopsis Biological Resource Center (ABRC) for providing seeds expressing P5R and P24Y.

## Author contributions

H.S. designed experiments and conception, and performed experiments and data analyses; Y.S. provided valuable suggestions and critical comments on this work; H.S. and C.S. wrote the manuscript. All authors read and approved the manuscript.

## References

Balla A, Tuymetova G, Barshishat M, Geiszt M, Balla T (2002) Characterization of Type II Phosphatidylinositol 4-Kinase Isoforms Reveals Association of the Enzymes with Endosomal Vesicular Compartments. J. Biol. Chem. 277: 20041– 20050

Barbosa ICR, Zourelidou M, Willige BC, Weller B, Schwechheimer C (2014) D6 PROTEIN KINASE Activates Auxin Transport-Dependent Growth and PIN-FORMED Phosphorylation at the Plasma Membrane. Dev. Cell. 29: 674–685

Barrett-Lennard EG, Dracup M (1988) A porous agar medium for improving the growth of plants under sterile conditions. Plant Soil 108: 294–298

Bleasdale JE, Bundy GL, Bunting S, Fitzpatrick FA, Huff RM, Sun FF, Pike JE (1989) Inhibition of phospholipase C dependent processes by U-73, 122. Adv Prostaglandin Thromboxane Leukot Res 19: 590–593.

Cho H-T, Cosgrove DJ (2002) Regulation of Root Hair Initiation and Expansin Gene Expression in Arabidopsis. Plant Cell 14: 3237–3253

Clough SJ, Bent AF (1998) Floral dip: a simplified method for Agrobacterium-mediated transformation of *Arabidopsis thaliana*: Floral dip transformation of Arabidopsis. Plant J. 16: 735–743

Drøbak BK (1992) The plant phosphoinositide system. Biochem. J. 288: 697–712

Durell WD (1941) THE EFFECT OF AERATION ON GROWTH OF THE TOMATO IN NUTRIENT SOLUTION. Plant Physiol. 16: 327–341

Fåhraeus G (1957) The Infection of Clover Root Hairs by Nodule Bacteria Studied by a Simple Glass Slide Technique. Microbiology. 16: 374–381

Kang B-H, Nielsen E, Preuss ML, Mastronarde D, Staehelin LA (2011) Electron Tomography of RabA4b- and PI-4Kβ1-Labeled Trans Golgi Network Compartments in Arabidopsis. Traffic. 12: 313–329

Ketelaar T, Anthony RG, Hussey PJ (2004) Green Fluorescent Protein-mTalin Causes Defects in Actin Organization and Cell Expansion in Arabidopsis and Inhibits Actin Depolymerizing Factor’s Actin Depolymerizing Activity in Vitro. Plant Physiol. 136: 3990–3998

Ketelaar T (2014) Live Cell Imaging of Arabidopsis Root Hairs. *In* V Žárský, F Cvrčková, eds, Plant Cell Morphogenesis. Humana Press, Totowa, NJ, pp 195– 199

Klein RR, Bourdon DM, Costales CL, Wagner CD, White WL, Williams JD, Hicks SN, Sondek J, Thakker DR (2011) Direct Activation of Human Phospholipase C by Its Well Known Inhibitor U73122. J. Biol. Chem. 286: 12407–12416

Koyama H, Toda T, Hara T (2001) Brief exposure to low-pH stress causes irreversible damage to the growing root in *Arabidopsis thaliana*: pectin–Ca interaction may play an important role in proton rhizotoxicity. J. Exp. Bot. 52: 361–368

Kusano H, Testerink C, Vermeer JEM, Tsuge T, Shimada H, Oka A, Munnik T, Aoyama T (2008) The *Arabidopsis* Phosphatidylinositol Phosphate 5-Kinase PIP5K3 Is a Key Regulator of Root Hair Tip Growth. Plant Cell. 20: 367–380

Loehwing WF (1934) PHYSIOLOGICAL ASPECTS OF THE EFFECT OF CONTINUOUS SOIL AERATION ON PLANT GROWTH. Plant Physiol. 9: 567–583

Marhava P, Aliaga Fandino AC, Koh SWH, Jelínková A, Kolb M, Janacek DP, Breda AS, Cattaneo P, Hammes UZ, Petrášek J, et al (2020) Plasma Membrane Domain Patterning and Self-Reinforcing Polarity in Arabidopsis. Dev. Cell. 52: 223–235.e5

Mueller-Roeber B, Pical C (2002) Inositol Phospholipid Metabolism in Arabidopsis. Characterized and Putative Isoforms of Inositol Phospholipid Kinase and Phosphoinositide-Specific Phospholipase C. Plant Physiol. 130: 22–46

Munnik T (2014) PI-PLC: Phosphoinositide-Phospholipase C in Plant Signaling. *In* X Wang, ed, Phospholipases in Plant Signaling. Springer Berlin Heidelberg, Berlin, Heidelberg, pp 27–54

Nakamura M, Claes AR, Grebe T, Hermkes R, Viotti C, Ikeda Y, Grebe M (2018) Auxin and ROP GTPase Signaling of Polar Nuclear Migration in Root Epidermal Hair Cells. Plant Physiol. 176: 378–391

Noack LC, Bayle V, Armengot L, Rozier F, Mamode-Cassim A, Stevens FD, Caillaud M-C, Munnik T, Mongrand S, Pleskot R, et al (2022) A nanodomain-anchored scaffolding complex is required for the function and localization of phosphatidylinositol 4-kinase alpha in plants. Plant Cell. 34: 302–332

Oancea E, Teruel MN, Quest AFG, Meyer T (1998) Green Fluorescent Protein (GFP)-tagged Cysteine-rich Domains from Protein Kinase C as Fluorescent Indicators for Diacylglycerol Signaling in Living Cells. J. Cell Biol. 140: 485–498

Okada K, Shimura Y (1990) Reversible Root Tip Rotation in *Arabidopsis* Seedlings Induced by Obstacle-Touching Stimulus. Science 250: 274–276

Ovečka M, Lang I, Baluška F, Ismail A, Illeš P, Lichtscheidl IK (2005) Endocytosis and vesicle trafficking during tip growth of root hairs. Protoplasma. 226: 39–54

Ovečka M, Berson T, Beck M, Derksen J, Šamaj J, Baluška F, Lichtscheidl IK (2010) Structural Sterols Are Involved in Both the Initiation and Tip Growth of Root Hairs in *Arabidopsis thaliana*. Plant Cell. 22: 2999–3019

Potters G, Pasternak TP, Guisez Y, Palme KJ, Jansen MAK (2007) Stress-induced morphogenic responses: growing out of trouble? Trends Plant Sci. 12: 98–105

Rounds CM, Lubeck E, Hepler PK, Winship LJ (2011) Propidium Iodide Competes with Ca^2+^ to Label Pectin in Pollen Tubes and Arabidopsis Root Hairs. Plant Physiol. 157: 175–187

Simon MLA, Platre MP, Assil S, Van Wijk R, Chen WY, Chory J, Dreux M, Munnik T, Jaillais Y (2014) A multi-colour/multi-affinity marker set to visualize phosphoinositide dynamics in Arabidopsis. Plant J. 77: 322–337

Stanislas T, Hüser A, Barbosa ICR, Kiefer CS, Brackmann K, Pietra S, Gustavsson A, Zourelidou M, Schwechheimer C, Grebe M (2015) Arabidopsis D6PK is a lipid domain-dependent mediator of root epidermal planar polarity. Nat. Plants 1: 15162

Staxén I, Pical C, Montgomery LT, Gray JE, Hetherington AM, McAinsh MR (1999) Abscisic acid induces oscillations in guard-cell cytosolic free calcium that involve phosphoinositide-specific phospholipase C. Proc. Natl. Acad. Sci. USA 96: 1779–1784

Stenzel I, Ischebeck T, König S, Hołubowska A, Sporysz M, Hause B, Heilmann I (2008) The Type B Phosphatidylinositol-4-Phosphate 5-Kinase 3 Is Essential for Root Hair Formation in *Arabidopsis thaliana*. Plant Cell. 20: 124–141

Takahashi H (1997) Hydrotropism: The current state of our knowledge. J. Plant Res. 110: 163–169

Thole JM, Vermeer JEM, Zhang Y, Gadella TWJ, Nielsen E (2008) *ROOT HAIR DEFECTIVE4* Encodes a Phosphatidylinositol-4-Phosphate Phosphatase Required for Proper Root Hair Development in *Arabidopsis thaliana*. Plant Cell. 20: 381–395

Van Leeuwen W, Vermeer JEM, Gadella TWJ, Munnik T (2007) Visualization of phosphatidylinositol 4,5-bisphosphate in the plasma membrane of suspension-cultured tobacco BY-2 cells and whole Arabidopsis seedlings. Plant J. 52: 1014– 1026

Vermeer JEM, Van Wijk R, Goedhart J, Geldner N, Chory J, Gadella TWJ, Munnik T (2017) In Vivo Imaging of Diacylglycerol at the Cytoplasmic Leaflet of Plant Membranes. Plant Cell Physiol. 58: 1196–1207

Verslues PE, Ober ES, Sharp RE (1998) Root Growth and Oxygen Relations at Low Water Potentials. Impact of Oxygen Availability in Polyethylene Glycol Solutions1. Plant Physiol. 116: 1403–1412

Watari M, Kato M, Blanc-Mathieu R, Tsuge T, Ogata H, Aoyama T (2022) Functional Differentiation among the *Arabidopsis* Phosphatidylinositol 4-Phosphate 5-Kinase Genes *PIP5K1*, PIP5K2 and PIP5K3. Plant Cell Physiol. 63: 635–648

Yanagisawa N, Kozgunova E, Grossmann G, Geitmann A, Higashiyama T (2021) Microfluidics-Based Bioassays and Imaging of Plant Cells. Plant Cell Physiol. 62: 1239–1250

Zhao J, Guo Y, Kosaihira A, Sakai K (2004) Rapid accumulation and metabolism of polyphosphoinositol and its possible role in phytoalexin biosynthesis in yeast elicitor-treated Cupressus lusitanica cell cultures. Planta. 219: 121–131

Zourelidou M, Müller I, Willige BC, Nill C, Jikumaru Y, Li H, Schwechheimer C (2009) The polarly localized D6 PROTEIN KINASE is required for efficient auxin transport in *Arabidopsis thaliana*. Development. 136: 627–636

