## Supplemental Figures and Tables for "Monitoring single root hairs using a micro-chambered hydroponics system MiCHy reveals a new mode of action of the aminosteroid U73122"

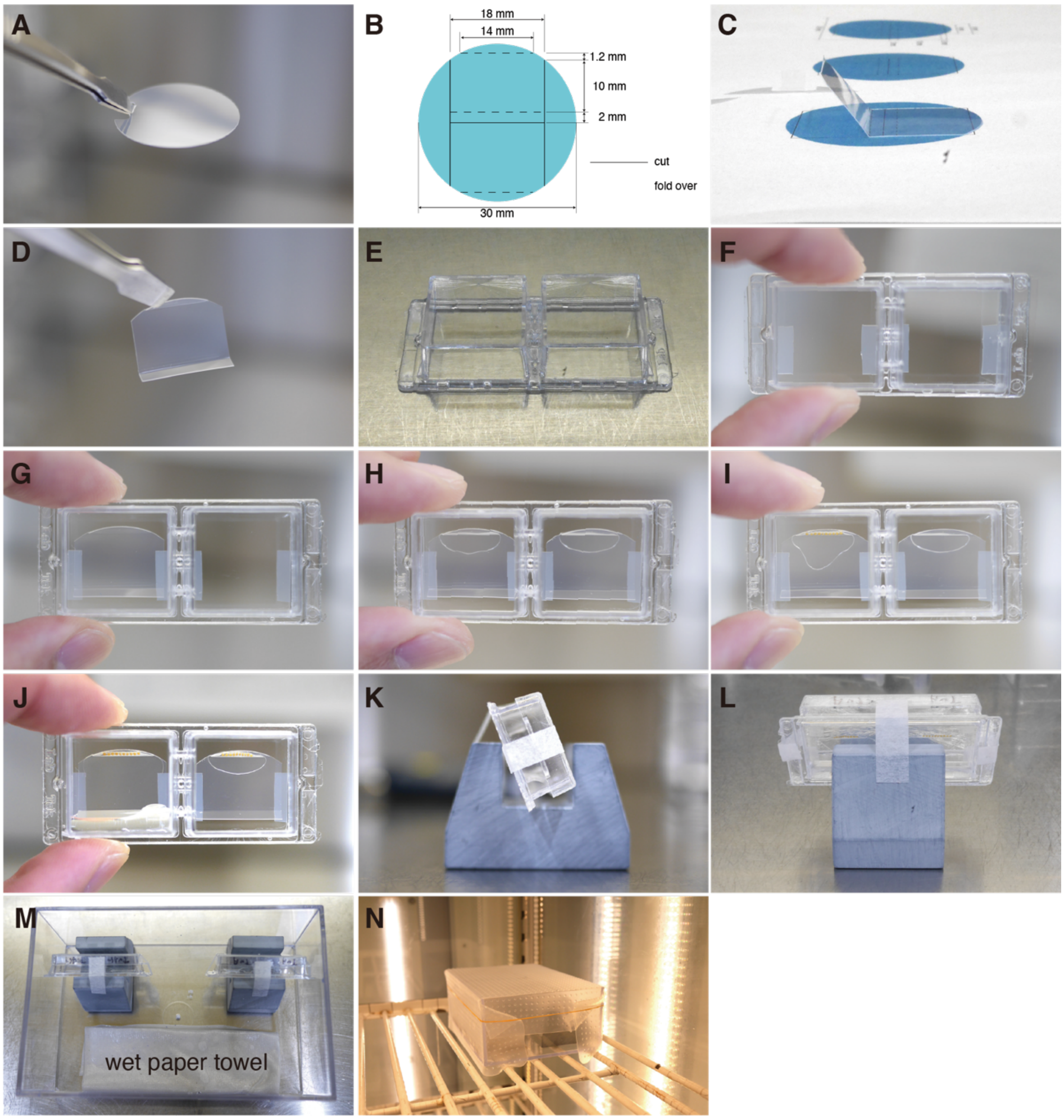


**Supplemental Fig. S1** Assembly of MiCHy. A sheet of the Cell Desk LF2 (**A**) is cut and folded according to the drawing (**B-D**). Slips of the silicone rubber sheet are attached to a chambered coverglass (**E** and **F**) and the prepared Cell Desk LF2 is layered on it (**G**). A minimal amount of melted 0.8% GTG-Agarose is injected into the thin space and solidified to form the bed for positioning Arabidopsis seeds (**H**). Arabidopsis seeds are placed on the bed (**I**) and then the liquid medium is poured into the bottom of the micro-chamber (**J**). The chamber is covered with the lid and held vertically (about 70˚) (**K** and **L**). The micro-chambers are held in a humid plastic box and covered with a silicon-rubber seal and kept in a growth cabinet for 5 to 6 days.

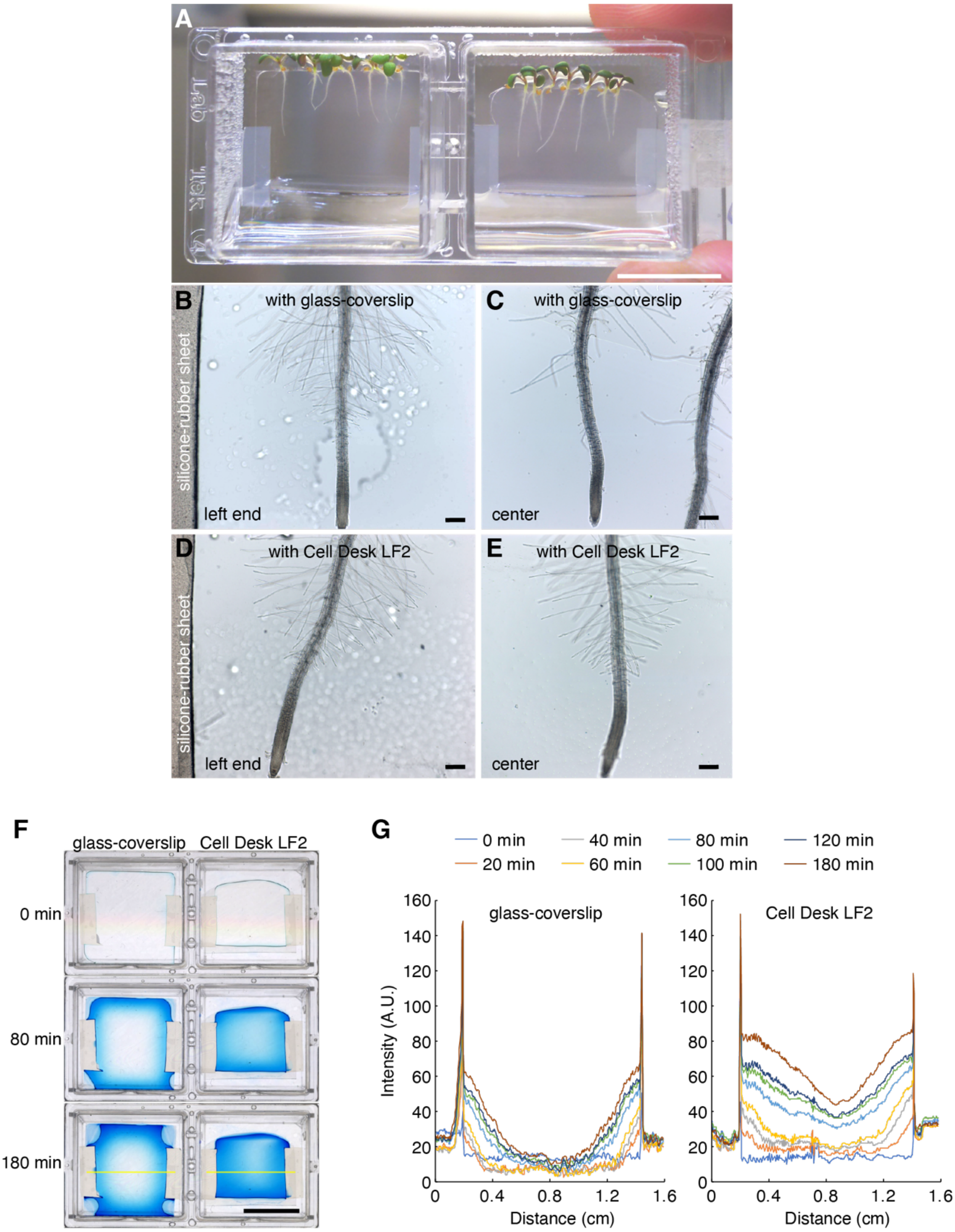


**Supplemental Fig. S2** Root growth in the micro-chambered hydroponics systems. **A)** The system using a glass coverslip in the left well and using Cell Desk LF2 in the right well. The white bar indicates 1 cm. **B**

and **C)** In the micro-chamber with the glass coverslip, roots and root hairs grow unevenly depending on their position within the chamber. **D** and **E)** In the micro-chamber with the Cell Desk LF2, the growth of roots and root hairs was uniform regardless of the position within the chamber. The bars indicate 200 μm. **F)** Detection of oxygen in the micro-chambers. Colorimetric oxygen detection was performed for 180 min. Blue-colored regions indicate the existence of oxygen. The black bar indicates 1 cm. **G)** Quantification of the colorimetric assay. The intensity profiles of the stained area on the line of interest for each micro-chamber in **F**) are presented.

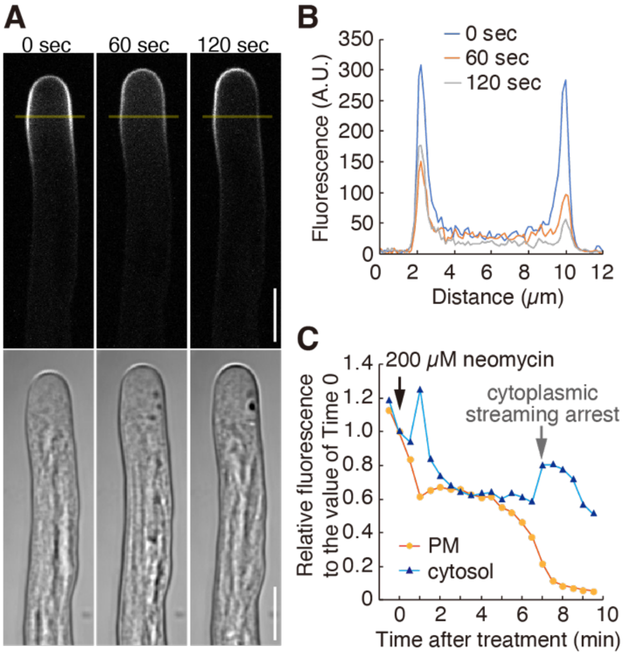


**Supplemental Fig. S3** Live-imaging of the PtdIns(4,5)$P_2$ biosensor responding to neomycin in MiCHy. **A)** Representative images of a root hair expressing CITRINE-PH$^{PLC}$ at 0, 60, and 120 sec after 200 µM neomycin treatment. White bars represent 10 µm. **B)** Distribution of fluorescence signals on the line of interest (yellow lines) in the images at 0, 60, and 120 sec after the treatment. **C)** Changes in fluorescence intensities on the plasma membrane (PM) or in the cytosol at the root hair tips. Quantification was performed as described in Supplemental Fig. S4, and the relative values to those at Time 0 are plotted. Arrows indicate the timing when 200 µM neomycin was applied. The experiments were repeated at least three times and the representative results are shown.

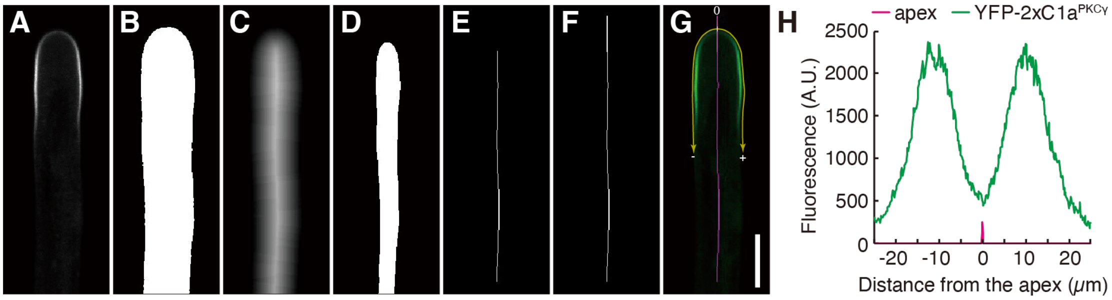


**Supplemental Fig. S4** Workflow for the image processing of root hairs. **A)** A raw image of a root hair expressing YFP-2xC1a$^{PKC\gamma}$ as an example. **B)** A binary image converted from the image A. **C.** A Euclidean distance map converted from the image B. **D)** A binary image converted from the image C. **E)** A skeletonized line converted from the image D. **F)** A vector of the edge part of the line E. is extended with 2-pixel width. **G)** A merged image of the images A and F. YFP-2xC1a$^{PKC\gamma}$ fluorescence and the vector line are shown in green and magenta, respectively. The intersection between the plasma membrane and the vector line is defined as an apex of the root hair. **H)** Fluorescence intensity profiles on the plasma membrane around the apex.

### Table S1. Methods using micro-chambers for live-imaging of roots and root hairs.

| | Micro-chamber | Medium state | Germinated (G) in or transplanted (T) into the chamber | Application | Species | References |
|---|---|---|---|---|---|---|
| Fåhraeus slide and derivertives | Thin medium sandwiched with a glass slide and coverslip | Solid | T | Imaging of root hair growth and the bacterial infection | Clover | Fåhraeus, 1957 |
| | Cell made from a glass slide and coverslip with glass spacers | Solid | T | Imaging of root hair growth and the bacterial infection | Legumes | Vincent, 1970 |
| | Cell made from a glass slide and coverslip, attached with silicone glue (also as spacer) | Liquid | T | Imaging of root hair growth and the bacterial infection | Clover | Bhuvaneswari and Solheim, 1985 |
| | Thin medium spread on a glass coverslip | Solid | G | Imaging of root hair growth and $Ca^{2+}$ with a chemical probe in root hairs | Arabidopsis | Wymer et al., 1997 |
| | | | | Imaging of actin in roots, including root hairs | Arabidopsis | Wang et al., 2007 |
| | | | | | | Dyachok et al., 2009 |
| | Thin space between a glass slide and coverslip | Liquid | G | Imaging of nuclei and actin in root hairs with chemical treatments | Arabidopsis | Ketelaar et al, 2002 |
| | | | T | Imaging of root hair growth with chemical treatments | Arabidopsis, *Triticum aestivum* | Ovecka et al., 2005 |
| | Thin medium spread on a coverglass, covered with gas-permealble film (biofoil) | Solid | G | Imaging of root hair growth and actin with chemical treatments | Arabidopsis | Ketelaar et al, 2004, 2014 |
| | Roots covered with a block of solid medium on a chambered-coverglass | Solid | T | Imaging of root growth with a vertical-stage microscopy | Arabidopsis | Von Wangenheim et al., 2017 |
| | | | | | | Rahni and Birnbaum, 2019 |
| | | | | | | Goh et al., 2022 |
| | Layered medium in a chambered coverglass | Solid | G | Imaging of $Ca^{2+}$, actin or nuclei in root hairs | Arabidopsis | Zhu et al., 2014 |
| | | | | | | Nakamura et al., 2018 |
| | Layered medium in a purpose-designed chambered coverglass (RoPod), assembled from a glasscoverslip and 3D-printed plastic ware | Solid | G | Imaging of root growth and chemical treatments | Arabidopsis | Guichard et al., 2021 |
| Custom perfusion chambers | Perfusion chamber made on a glass coverslip, chambered with silicone grease | Liquid | T | $Ca^{2+}$ imaging in root hairs | Alfalfa | Ehrhardt et al., 1996 |

| | | | | | |
|---|---|---|---|---|---|
| Perfusion chamber made on a glass coverslip, chambered with medical adhesive | Liquid | T | Glucose imaging with the FRET sensors in roots | Arabidopsis | Chaudhuri et al., 2011 |
| Perfusion chamber made with plexiglass and coverslips | Liquid | T | $Ca^{2+}$ imaging in roots | Arabidopsis | Zhu et al., 2014 |

### Table S2. Oligonucleotide and DNA sequences used in this study.

| # | Name | Description | DNA sequence |
|---|---|---|---|
| 1 | EcoRI-YFP_Fw | 1st and 2nd PCR for EcoRI-EYFP-AauI | 5’-AGAATTCATGGTGAGCAAGGGCGAGG -3’ |
| 2 | YFP_mutAauI_Rv | 1st PCR for EcoRI-EYFP-AauI fragment | 5’-AACTTGTGATGGGCTGCAGCTTATACAG -3’ underline, mutated AauI site in *YFP* ORF |
| 3 | YFP-AauI_Rv | 2nd PCR for EcoRI-EYFP-AauI fragment | 5’-TTTGTACAAACTTGTGATGGGCTG -3’ |
| 4 | EXP7pro_KpnI_Fw | KpnI-EXP7p-XhoI fragment | 5’-AAGGTACCGAGAGCGCCCGGTTTGAT -3’ |
| 5 | EXP7pro_XhoI_Rv | KpnI-EXP7p-XhoI fragment | 5’-TAACTCGAGTCTAGCCTCTTTTTCTTTATTCTTAGGG -3’ |
| 6 | 2xC1a$^{PKC\gamma}$ | Gene-synthesized fragment | 5’-GGGGACAAGTTTGTACAAAAAAGCAGGCTTCAGGCAAAAGGTTGTGCACGAGGTGAAGTCTCACAAGTTCACCGCTCGTTTCTTCAAGCAGCCTACCTTCTGCTCTCACTGCACCGATTTCATTTGGGGCATCGGAAAGCAGGGACTCCAATGTCAAGTTTGCTCTTTCGTGGTGCACAGAAGGTGCCACGAGTTCGTTACTTTTGAGTGTCCTGGTGGTGGAAGCGGAGGTGGTAGACAGAAAGTTGTTCATGAGGTCAAGAGCCATAAGTTCACGGCCAGATTTTTCAAGCAACCGACGTTCTGCAGCCACTGTACTGACTTCATCTGGGGAATTGGTAAGCAAGGGCTTCAGTGCCAGGTGTGCTCATTCGTTGTTCATAGACGTTGTCATGAGTTCGTGACCTTCGAGTGCCCTTAAGACCCAGCTTTCTTGTACAAAGTGGTCCCC-3’ |

### Supplemental References


**Bhuvaneswari TV, Solheim B** (1985) Root hair deformation in the white clover/*Rhizobium trifolii* symbiosis. *Physiol. Plant.* **63**: 25–34

**Chaudhuri B, Hormann F, Frommer WB** (2011) Dynamic imaging of glucose flux impedance using FRET sensors in wild-type Arabidopsis plants. *J. Exp. Bot.* **62**: 2411–2417

**Dyachok J, Yoo C-M, Palanichelvam K, Blancaflor EB** (2009) Sample Preparation for Fluorescence Imaging of the Cytoskeleton in Fixed and Living Plant Roots. *In* RH Gavin, ed, Cytoskeleton Methods and Protocols. Humana Press, Totowa, NJ, pp 157–169

**Ehrhardt DW, Wais R, Long SR** (1996) Calcium Spiking in Plant Root Hairs Responding to Rhizobium Nodulation Signals. *Cell.* **85**: 673–681

**Fåhraeus G** (1957) The Infection of Clover Root Hairs by Nodule Bacteria Studied by a Simple Glass Slide Technique. *Microbiology.* **16**: 374–381

**Goh T, Sakamoto K, Wang P, Kozono S, Ueno K, Miyashima S, Toyokura K, Fukaki H, Kang B-H, Nakajima K** (2022) Autophagy promotes organelle clearance and organized cell separation of living root cap cells in *Arabidopsis thaliana*. *Development.* **149**: dev200593

**Guichard M, Holla S, Wernerová D, Grossmann G, Minina EA** (2021) RoPod, a customizable toolkit for non-invasive root imaging, reveals cell type-specific dynamics of plant autophagy. Doi: 10.1101/2021.12.07.471480

**Ketelaar T, Faivre-Moskalenko C, Esseling JJ, De Ruijter NCA, Grierson CS, Dogterom M, Emons AMC** (2002) Positioning of Nuclei in Arabidopsis Root Hairs: An Actin-Regulated Process of Tip Growth. *Plant Cell.* **14**: 2941–2955

**Ketelaar T, Anthony RG, Hussey PJ** (2004) Green Fluorescent Protein-mTalin Causes Defects in Actin Organization and Cell Expansion in Arabidopsis and Inhibits Actin Depolymerizing Factor's Actin Depolymerizing Activity in Vitro. *Plant Physiol.* **136**: 3990–3998

**Ketelaar T** (2014) Live Cell Imaging of Arabidopsis Root Hairs. *In* V Žárský, F Cvrčková, eds, Plant Cell Morphogenesis. Humana Press, Totowa, NJ, pp 195–199

**Nakamura M, Claes AR, Grebe T, Hermkes R, Viotti C, Ikeda Y, Grebe M** (2018) Auxin and ROP GTPase Signaling of Polar Nuclear Migration in Root Epidermal Hair Cells. *Plant Physiol.* **176**: 378–391

**Ovečka M, Lang I, Baluška F, Ismail A, Illeš P, Lichtscheidl IK** (2005) Endocytosis and vesicle trafficking during tip growth of root hairs. *Protoplasma.* **226**: 39–54

**Rahni R, Birnbaum KD** (2019) Week-long imaging of cell divisions in the Arabidopsis root meristem. *Plant Methods.* **15**: 30

**Vincent, JM** (1970) A Manual for the Practical Study of the Root-Nodule Bacteria. IBP Handbook No. 15, Blackwell Scientific Publishers, 164p.

**Von Wangenheim D, Hauschild R, Fendrych M, Barone V, Benková E, Friml J** (2017) Live tracking of moving samples in confocal microscopy for vertically grown roots. *eLife.* **6**: e26792

**Wang Y, Yoo C, Blancaflor EB** (2007) Improved imaging of actin filaments in transgenic *Arabidopsis* plants expressing a green fluorescent protein fusion to the C- and N-termini of the fimbrin actin-binding domain 2. *New Phytol.* **177**: 525–536

**Wymer CL, Bibikova TN, Gilroy S** (1997) Cytoplasmic free calcium distributions during the development of root hairs of Arabidopsis thaliana. *Plant J.* **12**: 427–439

**Zhu X, Taylor A, Zhang S, Zhang D, Feng Y, Liang G, Zhu J-K** (2014) Measuring Spatial and Temporal $Ca^{2+}$ Signals in Arabidopsis Plants. *JoVE.* 51945

**Supplemental Movies**

**Supplemental Movie S1** Diffusion of 0.25% Evans blue dye in the agar medium. The dye was applied into the micro-chamber based on Nakamura et al., 2008.

**Supplemental Movie S2** Detection of oxygen diffusion in micro-chambers.

**Supplemental Movie S3** Chemical treatment procedure in MiCHy.

**Supplemental Movie S4** Wash-out procedure in MiCHy.

**Supplemental Movie S5** FM4-64X staining of a main root in MiCHy. FM4-64X fluorescence and bright field images in time-lapse are presented in the left and right panels, respectively. The images are maximum intensity projections of 250-μm thickness.

**Supplemental Movie S6** PI staining of a main root in MiCHy. PI fluorescence and bright field images in time-lapse are presented in the left and right panels, respectively. The images are maximum intensity projections of 195-μm thickness.

**Supplemental Movie S7** Filipin III staining of a growing root hair in MiCHy. Filipin III fluorescence and bright field images in time-lapse are presented in the upper and bottom panels, respectively. The bar indicates 10 μm.

**Supplemental Movie S8** BFA treatment and its wash-out for a YFP-D6PK-expressing root in MiCHy. YFP fluorescence and bright field images in time-lapse are presented in the left and right panels, respectively. The images are maximum intensity projections of 50-μm thickness.

**Supplemental Movie S9** Treatment with 0.1% DMSO for a root hair expressing PtdIns(4)P biosensor 2xCherry-$PH^{FAPP1}$ and PtdIns(4,5)$P_2$ biosensor CITRINE-$PH^{PLC}$ in MiCHy. The images are sum slices intensity projection of 0.5-μm thickness. The bar indicates 10 μm.

**Supplemental Movie S10** Treatment with 30 μM PAO for a root hair expressing PtdIns(4)P biosensor 2xCherry-$PH^{FAPP1}$ and PtdIns(4,5)$P_2$ biosensor CITRINE-2x$PH^{PLC}$ in MiCHy. The images are sum slices intensity projection of 0.5-μm thickness. The bar indicates 10 μm.

**Supplemental Movie S11** Treatment with 20 μM U73122 for a root hair expressing PtdIns(4)P biosensor 2xCherry-PH$^{FAPP1}$ and PtdIns(4,5)$P_2$ biosensor CITRINE-2xPH$^{PLC}$ in MiCHy. The images are sum slices intensity projection of 0.5-μm thickness. The bar indicates 10 μm.

**Supplemental Movie S12** Treatment with 5 μM U73122 for a root hair expressing PtdIns(4,5)$P_2$ biosensor CITRINE-2xPH$^{PLC}$ in MiCHy. YFP fluorescence and bright field images in time-lapse are presented in the upper and bottom panels, respectively. The images are sum slices intensity projection of 0.5-μm thickness. The bar indicates 10 μm.

**Supplemental Movie S13** Treatment with 200 μM neomycin for a root hair expressing PtdIns(4,5)$P_2$ biosensor CITRINE-2xPH$^{PLC}$ in MiCHy. YFP fluorescence and bright field images in time-lapse are presented in the upper and bottom panels, respectively. The images are sum slices intensity projection of 1-μm thickness. The bar indicates 10 μm.

**Supplemental Movie S14** Treatment with 0.1% DMSO, 5 μM U73122, or 200 μM neomycin for root hairs expressing DAG biosensor YFP-2xC1a$^{PKC\gamma}$ in MiCHy. YFP fluorescence and bright field images in time-lapse are presented in the left and right panels, respectively. The images are sum slices intensity projection of 1-μm thickness. The bar indicates 10 μm.

**Supplemental Movie S15** Treatments with 0.1% DMSO, 5 μM U73122, or 200 μM neomycin for root hairs expressing PIP5K3-YFP in MiCHy. YFP fluorescence and bright field images in time-lapse are presented in the left and right panels, respectively. The images are sum slices intensity projection of 0.6-μm thickness. The bar indicates 10 μm.
